# Seasonal niche overlap of diverse facultative antagonistic bacteria of diatoms in a productive coastal ecosystem

**DOI:** 10.1101/2024.06.05.594869

**Authors:** Laura Branscombe, Cordelia Roberts, Claire Widdicombe, Courtney Swink, William H. Wilson, Michael Cunliffe, Katherine Helliwell

## Abstract

Biotic interactions between microbes underpin marine ecosystem health, governing the flux of carbon and other nutrients in the ocean. However, studying aquatic microbial interactions is challenging. Model systems can provide in depth understanding of the mechanisms driving such associations. Yet, insights of the prevalence and co-occurrence dynamics of laboratory model systems in natural ecosystems remain limited. By leveraging 16S and 18S metabarcoding combined with phylogenetic analysis, we assessed the environmental presence of facultative bacterial pathogens of one of the most globally abundant phytoplankton groups, the diatoms. Sampling microbial assemblages in a productive coastal ecosystem over the course of an annual cycle, we detected multiple algicidal bacteria that frequently exhibited overlapping co-occurrences. Together, these bacteria positively correlated with members of the potentially toxic genus *Pseudo-nitzchia*, as well as temperature. Our study indicates that antagonistic bacteria occupy shared temporal niches and demonstrates the need to consider their cumulative impacts on diatom population health, including in future ocean conditions.

## Introduction

As one of the most successful phototrophic groups in the global ocean (Malviya *et al*. 2016), diatoms contribute approximately 40% of marine primary productivity and drive major nutrient (e.g. nitrogen and silica) cycles (Nelson *et al*. 1995; Kamp *et al*. 2016). Diatoms are an important source of organic carbon for higher and lower trophic organisms, fuelling marine ecosystems through primary productivity (Buchan *et al*. 2014). Seasonal diatom blooms are significant events that support an enormous diversity of microbes, including heterotrophic bacteria, which utilise the substantial amounts of organic carbon that result from a diatom bloom. Early studies demonstrated that around half of algal derived primary production is consumed by marine bacteria (Cole, Findlay and Pace 1988; Ducklow *et al*. 1993). The remaining carbon is either utilised by other heterotrophic organisms, or sequestered in marine sediments when diatom aggregates sink to the sea floor (Jiao *et al*. 2010; Giering *et al*. 2014). Understanding factors driving diatom success and bloom dynamics in marine environments is thus of paramount importance for understanding microbial community dynamics and global biogeochemical cycles.

Diatoms engage in multifaceted interactions with bacteria. Bacteria can produce a range of diatom growth promoting compounds, from vitamins (Haines and Guillard 1974; Croft *et al*. 2005; Durham *et al*. 2017) and iron-chelating siderophores (Amin *et al*. 2009), to hormones such as indole acetic acid (Amin *et al*. 2015). In turn, bacteria benefit from diatom-derived dissolved organic matter, including sugars and cofactors as well as signalling molecules (Shibl *et al*. 2020). These metabolites vary between diatom hosts, and also according to diatom physiology (e.g. culture age) (Brisson *et al*. 2023). Moreover, they can have differential impacts on bacterial growth with ‘functional guilds’ of bacteria able to utilise or remineralise distinct classes of molecules to different extents (Mayali *et al*. 2023).

Competitive and antagonistic interactions between diatoms and bacteria are also widely cited. Originally isolated from a culture of the harmful diatom *Pseudo-nitzchia multiseries* (Amin *et al*. 2015; Van Tol, Amin and Armbrust 2017), the *Gammaproteobacter Croceibacter atlanticus* inhibits diatom growth and disrupts cell cycle progression (Van Tol, Amin and Armbrust 2017; Bartolek *et al*. 2022). Similarly, another algicidal flavobacterium, *Kordia algicida,* secretes proteases that cause diatom cell death (Paul and Pohnert 2011; Syhapanha *et al*. 2023). Subsequent mesocosm experiments have shown *K. algicida* caused shifts in community composition of natural phytoplankton populations, including a rapid decline of the diatom *Chaetoceros socialis* followed by a sharp increase in the haptophyte *Phaeocystis* (Bigalke and Pohnert 2019). However, despite the clear implications of algicidal bacteria to diatom health and bloom succession in nature, there remains a significant chasm between studies of laboratory model systems that can provide crucial insights of the biology underlying such interactions and understanding of the presence and significance of such algicidal bacteria in nature. While co-occurrence network analysis has proven informative in inferring negative and positive associations between diatoms and other microbes in aquatic ecosystems (Vincent and Bowler 2020; Siebers *et al*. 2024), further work is necessary to deduce the nature of such putative interactions. The aim of this study is thus to take a reverse approach to quantify the prevalence and co-occurrence dynamics over time of bacterial taxa known through laboratory study to exert substantial negative impacts on diatom growth and fitness.

Our recent work employing plaque-assay sampling to isolate a library of antagonistic bacteria of diatoms from L4 Station in the Western English Channel (WEC) identified multiple bacteria capable of causing severe detrimental effects on diatom growth (Branscombe *et al*. 2024). Laboratory characterisation of eight of these strains revealed that in all cases, these effects were facultative, activated by pre-exposure to diatom necromass. The diatom-attaching Roseobacter species *Ponticoccus alexandrii,* was particularly persistent in this region being isolated on multiple independent occasions. *P. alexandrii* caused species-specific impacts on diatom growth and viability, but either had no, or far reduced, effects against other algal (haptophyte and dinoflagellate) taxa. The observed taxa-specific effects further highlight the potential of algicidal bacteria to shape phytoplankton species succession. Indeed, notably, we identified peaks in plaque enumeration suggesting increased bacterial pathogen load during senescence of a winter bloom of the centric diatom *Coscinodiscus*. Moreover, detection of metabarcodes phylogenetically similar to several bacterial antagonists isolated from the WEC in marine ecosystems globally (Branscombe *et al*. 2024), indicates the broader biogeography of such bacteria.

By virtue of being isolated from the WEC that is home to one of the longest running and well-studied oceanographic time series globally, the Western Channel Observatory, the strains described offer ‘environmental tractability’. The close proximity of L4 Station off the coast of Plymouth lends itself to frequent sampling of diatom and bacterial assemblages (Gilbert *et al*. 2009; Caporaso *et al*. 2012; Taylor *et al*. 2014; Taylor and Cunliffe 2016) and is underpinned by extensive long-term metadata of phytoplankton diversity and physicochemical parameters (McEvoy *et al*. 2023). Exploiting these attributes combined with our previous knowledge gained from isolating facultative antagonistic bacteria from this region (Branscombe *et al*. 2024), we aimed to assess the presence of such bacteria (as well as additional algicidal bacteria reported in the literature) to study their co-occurrence with phytoplankton in this productive coastal ecosystem where diatoms frequently bloom, using 16S and 18S metabarcoding.

## Materials and Methods

### Seawater sample collection and processing

Seawater samples for metabarcoding were collected from Station L4 (50° 15.00′ N, 4° 13.02′ W) by RV Sepia (Marine Biological Association, UK). Samples were taken bi-monthly (where possible, with the exception of March and April 2021, where samples were collected weekly for greater temporal resolution during the spring months) for 13 months (**Table S1**) from a depth of 5 m. For each sampling point, four 1 L seawater replicates were filtered through a 0.22 µm cellulose nitrate filter paper membrane (Cole-Palmer, UK), which was immediately stored in a DNA/RNA shield reagent (Zymo Research, USA) at –80 °C until DNA extraction.

### DNA extraction and metabarcoding

Samples were thawed on ice and DNA was extracted from the filters using a ZymoBIOMICS DNA Miniprep kit (Zymo Research, USA). Thawed filters were cut into smaller sections and extractions were carried out according to the manufacturer’s instructions, with the addition of a mechanical bead-beating step. For eukaryotic community analysis, 18S rRNA gene amplification of the V9 region was conducted via PCR using the primers 1391F (GTACACACCGCCCGTC) and EukB (TGATCCTTCTGCAGGTTCACCTAC) (Lane *et al*. 1985; Medlin *et al*. 1988), an initial denaturation step at 95 °C for three minutes, followed by 35 cycles of i) denaturation at 95 °C for 45 seconds, ii) annealing at 57 °C for one minute, and iii) extension at 72 °C for one minute and 30 seconds. Final extension was carried out at 72 °C for ten minutes. For prokaryotic community analysis, 16S rRNA gene amplification of the V4 region was conducted via PCR using the primers 515F (GTGCCAGCMGCCGCGGTAA) and 806R (GGACTACHVGGGTWTCTAAT) (Caporaso *et al*. 2011). PCR cycle conditions were as follows: an initial denaturation step at 94 °C for two minutes, followed by 35 cycles of i) denaturation at 94 °C for 45 seconds, ii) annealing at 50 °C for one minute, and iii) extension at 72 °C for one minute and 30 seconds. PCR products were then analysed via gel electrophoresis by running on a 1% agarose gel at 110 volts for 60 minutes to check for amplification. Final DNA concentrations were quantified using a NanoDrop spectrophotometer (ThermoFisher). Both 16S and 18S rRNA gene amplicons were sequenced on the Illumina MiSeq platform.

### Data processing of amplicon sequence data

Amplicon sequence data was processed in R Studio (R Core Team, 2019) following the DADA2 pipeline (Callahan *et al*. 2021). 16S and 18S samples were processed independently of each other following the same pipeline: primers and low-quality sequences were removed by filtering and trimming demultiplexed reads, before merging paired ends to produce full denoised sequences. Chimeric sequences were subsequently removed prior to taxonomic assignment of amplicon sequence variants (ASVs) using the PR^2^ database (release 4.140) (Guillou *et al*. 2013) for 18S sequences, and the SILVA database version 138.1, (Quast *et al*. 2013) for 16S sequences. Chloroplast and mitochondrial sequences were removed from both data sets, and non-eukaryote sequences were removed from the 18S data set.

The phyloseq package (McMurdie and Holmes 2013) was then used to create a phyloseq object by combining ASV tables, taxonomic assignments and sample metadata. Read depths of each sample were inspected, and sequences were rarefied to a depth of 10,443 reads per sample for 16S sequences, and 10,716 for 18S sequences. To avoid losing diversity of samples with higher read depths, four samples with comparatively low read depths (< 10,000 reads) were removed from the 16S dataset (**Table S2**).

### Investigating the occurrence of bacterial antagonists within the WEC

To assess the abundance and seasonal trends of bacterial antagonists, separate 16S BLAST databases were created using a non-rarified 16S amplicon dataset in Geneious Prime (Geneious Prime 2023.0.1, https://www.geneious.com). Full-length 16S rRNA gene sequences (**Dataset S1**) (Sohn *et al*. 1978; Wang *et al*. 2016; Van Tol, Amin and Armbrust 2017; Branscombe *et al*. 2024) were queried against the 16S ASV database, and ASVs with a percentage pairwise identity above 97% (query cover 100%) were collated. Taxonomic assignment of each ASV was subsequently validated by constructing Maximum Likelihood trees.

### Verification of taxonomic assignment of ASV hits via construction of Maximum Likelihood phylogenetic trees

Maximum Likelihood trees were constructed in Geneious Prime (Geneious Prime 2023.0.1, https://www.geneious.com) to verify the taxonomic assignment of ASV hits obtained through local BLAST searches of the bacterial amplicon sequence dataset. Maximum Likelihood trees were constructed using the full-length 16S rRNA bacterial query sequence, the V4 region of each nearest ASV hit, and full-length 16S rRNA sequences of type strains of closely related species and genera obtained from NCBI and the Ribosomal Database Project (Cole *et al*. 2014; Sayers *et al*. 2022). Sequences were aligned using a Multiple Alignment Fast Fourier Transform (MAFFT) tool (Katoh *et al*. 2002), and alignments were then manually trimmed to standardise sequence lengths, before using the General-Time-Reversable (GTR) substitution model with 1000 bootstraps to construct Maximum Likelihood trees.

### Measurement of phytoplankton and bacterial abundances as well as environmental parameters at L4 Station

In addition to amplicon sequence analysis, phytoplankton and bacterioplankton abundance (cells/ml) as well as physicochemical environmental parameters were monitored via the Western Channel Observatory (McEvoy *et al*. 2023), as follows. For total phytoplankton counts (of cells >2 µm), seawater samples were collected bi-weekly (where possible) from Station L4 from a depth of 10 m. Samples were fixed in 2% Lugol’s iodine solution for enumeration by light microscopy according to the Utermohl counting technique (Utermohl 1958). Organisms were classified into taxonomic groups (i.e., diatoms, coccolithophores, dinoflagellates, ciliates, etc.) and identified to species level where possible. Biomass was calculated by converting biovolume data to carbon (µg C l^-1^) following the Menden-Deuer and Lessard formula (Menden-Deuer and Lessard 2000). Bacterial abundance was quantified using flow cytometry with protocols outlined in (Tarran and Bruun 2015). Nutrient concentrations (nitrate + nitrite, silicate and phosphate) were monitored according to (Woodward and Rees 2001). Temperature, salinity and photosynthetically active radiation were recorded via autonomous Conductivity Temperature Depth (CTD)-sensors.

### Statistical analyses

Correlations between ASVs phylogenetically similar to facultative antagonistic bacteria and abiotic (nutrients, alongside temperature, salinity and photosynthetically active radiation (PAR)) as well as biotic (phytoplankton counts) metadata were determined using two-way Pearson’s multivariate analysis in GraphPad Prism 10.1.1.

## Results

### Overview of diatom bloom and bacterial abundance dynamics at L4 Station

Phytoplankton and bacterial count abundance data collated from the Western Channel Observatory revealed overarching trends in the abundance and seasonal patterns of phytoplankton and bacterial abundance between March 2021 to January 2022 (**Figure 1A**). Diatoms were by far the most abundant eukaryotic phytoplankton group throughout the sampling window, with the exception of the third week of April 2021 and also during early Oct 2021 when dinoflagellates and coccolithophorids exceeded diatom cell counts, respectively (**Figure S1A**). Peaks in total diatom biomass occurred as expected during the spring (with two major peaks in April and May 2021) (**Figure 1A**, left hand axis). A summer bloom was also apparent in mid-August, with subsequent smaller peaks in diatom biomass detected during autumn (September to October) and to a lesser degree winter (January 2022). Total bacterial abundance (cell counts), as monitored via flow cytometry (**Figure 1A**, right hand axis), was maximal in the third week of June 2021 (with 3.8 x 10^6^ cells per ml), following the largest diatom bloom observed during our sampling period that occurred in the fourth week of May 2021. Bacterial abundances subsequently declined gradually over the summer months to below 1 x 10^6^ cells per ml (at the start of August 2021), with a modest increase later that month coinciding with the summer diatom bloom followed by another peak in November. Bacterial cell counts subsequently remained low thereafter. Metadata of abiotic environmental variables (**Figure S1B-D**) indicated depletion of inorganic nutrients (nitrite + nitrate, phosphate and silicate) towards the end of April (after the first peak in diatom biomass), and these nutrients remained low until mid-August 2021 (**Figure 1A**; **Figure S1B**). Peaks in temperature were observed in July **(Figure S1C**).

**Figure 1.**
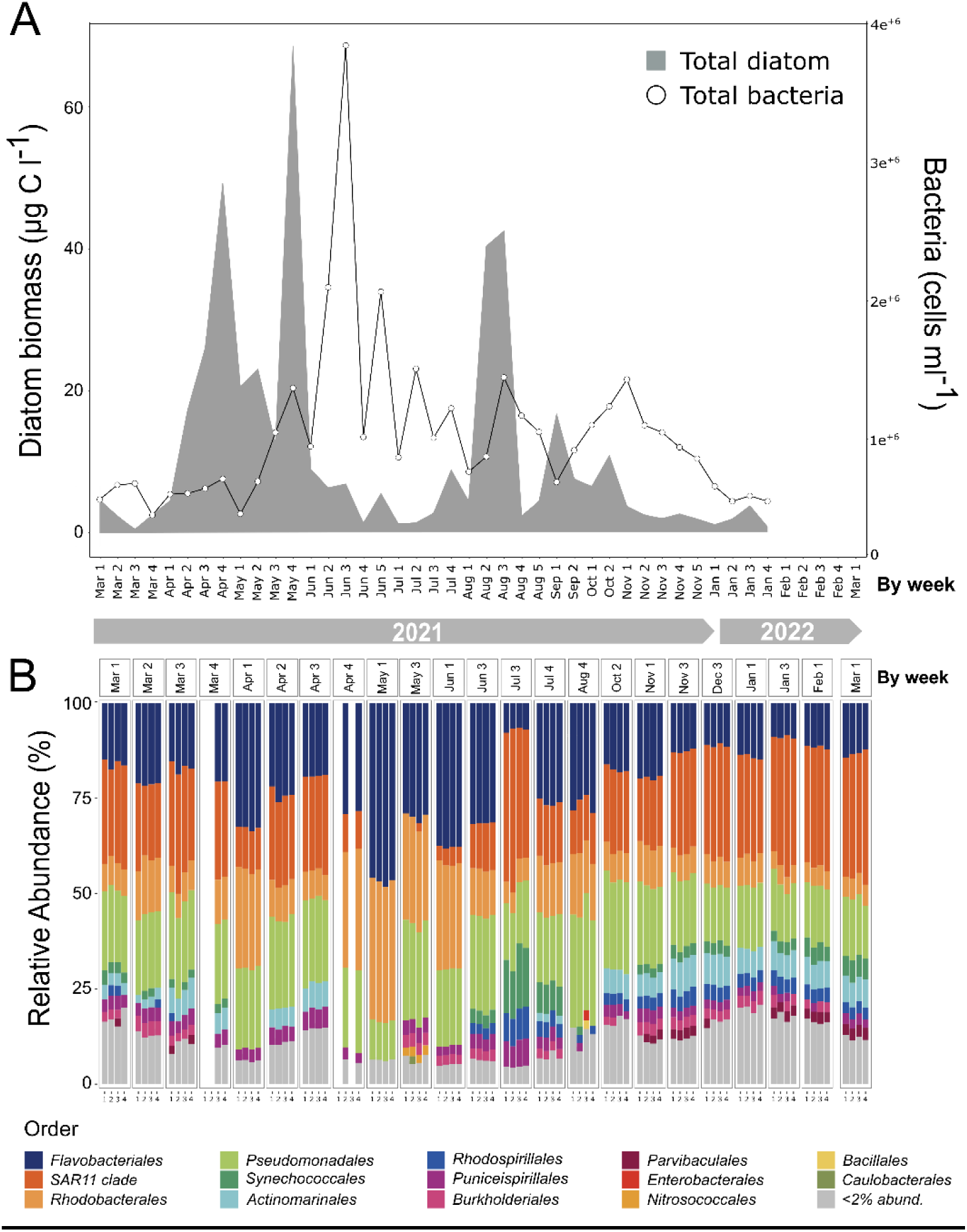
Diatom and bacterial abundance dynamics at L4 Station over an annual station. **A.** Biomass (µg C l^-1^) of total diatoms between March 2021 and January 2022, indicated by sampling week. Also included is total bacterial abundance (cells ml^-1^), demonstrating peak abundance upon the termination of a diatom bloom in May 2021. Nb. Plankton abundance data beyond February 2022 was not available from the Western Channel Observatory at the time of writing this manuscript. **B.** Percentage relative abundance of amplicon sequencing variants (ASVs) belonging to major bacterial orders (March 2021 – March 2022). Each sampling point contains four replicates. Taxa that constituted less than 2% of the total ASVs are grouped into the <2% abundance category. Samples (replicates) with read depths <10,000 were removed to avoid losing diversity of samples with higher read depths in the rarefication step of the sequencing pipeline.

### Characterisation of the bacterial community using 16S rRNA gene metabarcode analysis

The diversity and relative abundance of specific bacterial taxa at L4 Station during the sampling period was characterised further by 16S rRNA gene metabarcoding. Analysis of 16S rRNA gene amplicon sequence data revealed that the bacterial community was consistently largely dominated by four groups, *Flavobacteriales*, *SAR11* clade, *Rhodobacterales* and *Pseudomonadales* (**Figure 1B**). These four groups remained the most abundant taxa throughout the entire annual cycle, except for the May 2021 sampling points, where the *SAR11* clade decreased substantially to less than 2% of the total relative abundance of the prokaryotic community. The decrease in *SAR11* clade abundance in the first week of May was concurrent with increases in the relative abundance of the *Flavobacteriales* and the *Rhodobacterales* and coincided with subsidence of the first peak of the spring diatom bloom (i.e. the 1^st^ and 2^nd^ weeks of May 2021, **Figure 1A**). However, maximum bacterial numbers were not observed until the 3^rd^ week of June (**Figure 1A**), after the second diatom bloom peak (in the last week of May 2021). *Pseudomonadales* were also seasonally persistent throughout the entire sampling period but remained more stable across the annual cycle compared to the *Flavobacteriales*, *Rhodobacterales* and *SAR11*, often constituting approximately 20% of total prokaryotic reads (with the exception of the first week of May 2021, where relative abundance decreased).

### Detection of ASVs for laboratory characterised antagonistic bacteria of diatoms at L4 Station

We sought to determine the presence of bacterial antagonists of diatoms in our 16S rRNA gene metabarcoding data. We searched our non-rarefied prokaryote amplicon sequence data for facultative antagonistic bacterial taxa we have previously isolated from the WEC that confer robust growth inhibitory effects against diatoms in laboratory culture (Branscombe *et al*. 2024). We queried the 16S rRNA gene sequences of *P. alexandrii*, *T. lohafexi*, *M. spongiicola*, *H. titanicae*, *M. adhaerens*, *M. idriensis*, and *V. diazotrophicus* (**Dataset S1**). Based on sequence similarity we identified putative positive hits for all bacteria, including three ASVs for *P. alexandrii*, two for *M. adhaerens*, *M. spongiicola*, *M. idriensis* each and also one hit for *T. lohafexi*, *V. diazotrophicus* and *H. titanicae,* respectively (**Table S3**). Maximum Likelihood trees were constructed to further scrutinise the taxonomic assignment of each hit. This analysis identified with robust bootstrap support an ASV for *P. alexandrii* (**Figure S2A**; asv_1208), *T. lohafexi* (or close relative *T. lucentensis*) (**Figure S2B**; asv_1398), as well as *M. adhaerens/M. flavimaris* (**Figure S2C**; asv_989). Additionally, we identified an ASV each for *M. idriensis* (**Figure S3A**; asv_1597) and *V. diazotrophicus* (**Figure S3B**; asv_1535). The ASV hits for *M. spongiicola* did not group together with that of our previous *M. spongiicola* isolate (nor with the reference sequence for this species) (**Figure S3C**). However, sequences were instead found to be closely related to *Maribacter dokdonensis* (asv_3491 and asv_3288) previously isolated from the WEC with confirmed algicidal activity towards diatoms (Wang *et al*. 2016; Branscombe *et al*. 2024). Finally, the ASV hit for the *H. titanicae* query sequence (asv_774) clustered most closely with *Halomonas sulfidaeris* (**Figure S4A**).

In addition to examining bacterial antagonists isolated previously from the WEC (Wang *et al*. 2016; Branscombe *et al*. 2024), we searched for other bacteria known to be algicidal towards diatoms from literature reports, including *C. atlanticus* (Van Tol, Amin and Armbrust 2017), *K. algicida* (Paul and Pohnert 2011) and *Alteromonas macleodii* (Cai *et al*. 2023). Two ASVs with high sequence similarity to *C. atlanticus* were identified (**Table S3**; asv_1181 and asv_2324), both of which were later verified to be taxonomically similar to *C. atlanticus* by Maximum Likelihood tree construction (**Figure S4B**). Finally, querying *A. macleodii* against the dataset also returned two ASVs with high % sequence similarity (**Table S3**). Maximum Likelihood tree construction revealed one of the ASVs (asv_1961) to cluster robustly with another algicidal strain *Alteromonas* sp. PML-EC1, previously isolated from the WEC (Wang *et al*. 2016), while the second ASV (asv_934) clustered more closely with *Alteromonas tagae* (**Figure S4C**), despite having a percentage pairwise identity of 100% with the *A. macleodii* query sequence. A query search of *K. algicida* against the dataset did not return any ASVs with sequence similarity above 70%.

### Seasonal trends in the abundance of verified facultative bacterial antagonists and phytoplankton community composition

Each confirmed ASV described above was subsequently monitored throughout the amplicon sequencing sampling period. We detected ASVs for antagonistic bacterial taxa in multiple temporally distinct samples from spring to the following winter (in March, April, June, July, August, October as well as November), including in 11 of the 23 sampling points of this time series (**Figure 2A**). On several occasions we detected multiple bacterial antagonists co-occurring at the same time, most notably towards the end of June and August 2021. The latter time-point coincided with the demise of the summer diatom bloom (**Figure 2A**). ASVs observed in August 2021 included *Halomonas* sp., *M. adhaerens*, *T. lohafexi*, and *Alteromonas* sp. (asv_1961). By comparison, in late June we detected two *C. atlanticus* ASVs as well as *P. alexandrii* and *M. dokdonensis* (**Figure 2A**). The first detection of *C. atlanticus* (asv_1181) was in the first week of June 2021, just as the May bloom was subsiding. Additionally, ASVs for *Alteromonas* sp. (asv_1961) and *V. diazotrophicus* were detected at the end of July. We also identified several ASV hits in early spring, including for *P. alexandrii* (late March) as well as for *Halomonas* sp. (March and April sampling points). *M. idriensis* was identified in November (**Figure 2A**).

**Figure 2.**
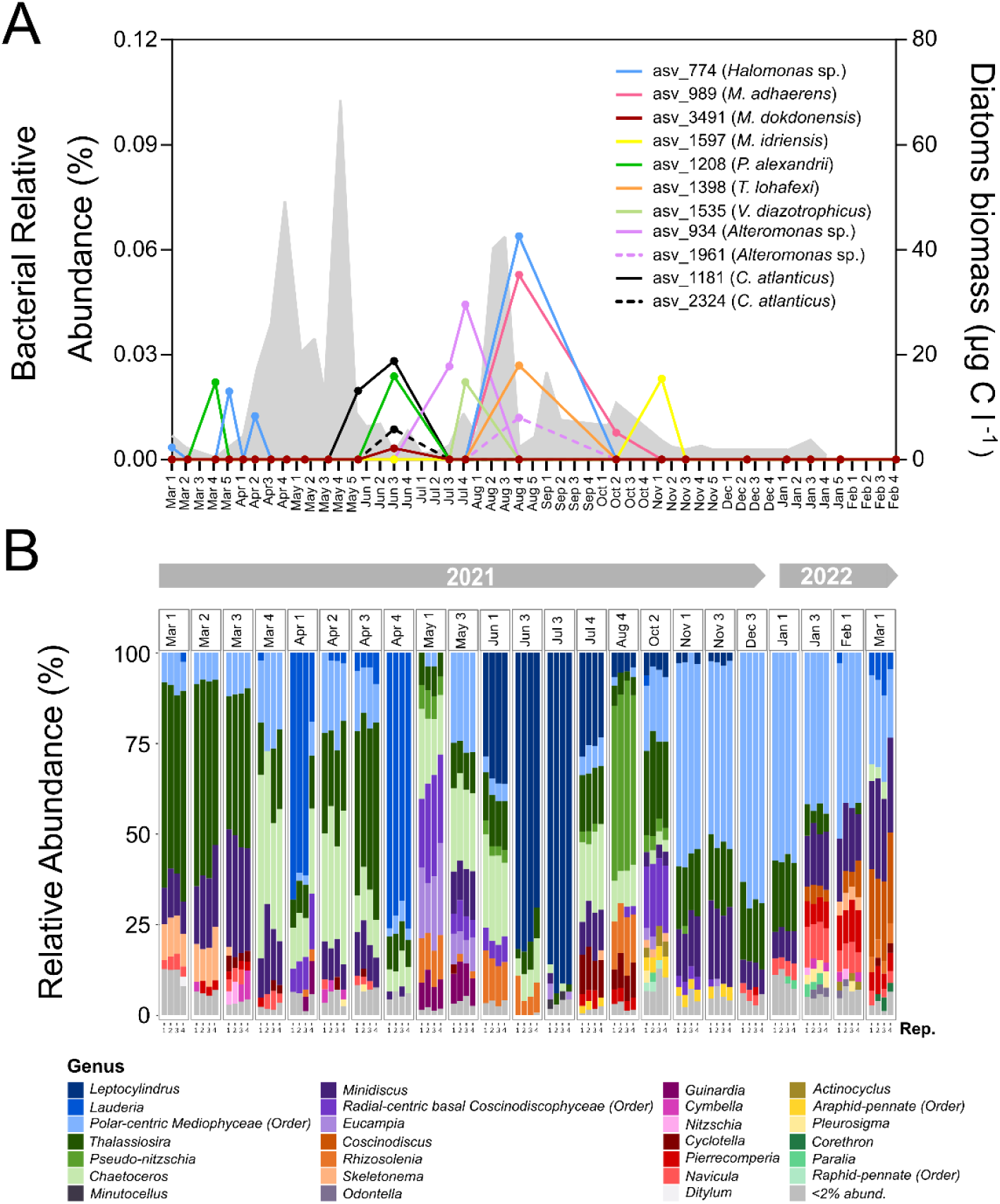
Co-occurrence dynamics of antagonistic bacteria and diatom assemblages at L4 Station. **A.** Percentage relative abundance (calculated as the sum of reads detected in each four replicates collected per sampling point as a percentage of the total number of bacterial ASVs across the four replicates) of ASVs phylogenetically related to query sequences of bacterial antagonists with confirmed facultative antagonistic activity. Total diatom biomass (µg C l^-1^, grey, right axis) over an annual cycle, labelled by sampling week) is also shown. **B.** Relative abundance (%) of major diatom genera over the same sampling period. Each sampling point contains four replicates. Taxa that constituted less than 2% of the total ASVs are grouped into the <2% abundance category.

To determine which specific diatom taxa were present and co-occurring with bacteria identified, we conducted 18S rRNA gene amplicon sequencing of the eukaryotic community. This confirmed that diatoms constituted a numerically important component of the phytoplankton community year-round (yellow, **Figure S5**), albeit dinoflagellates exhibited the greatest relative abundance in the amplicon sequencing data, in contrast to the Western Channel Observatory cell counts (**Figure S1A**) (most likely an artefact of the high copy number of 18S rRNA genes in dinoflagellates (Ruvindy et al. 2023). Relative abundance of major diatom genera (*Leptocylindrus*, *Thalassiosira*, *Chaetoceros*, and *Lauderia*) derived from 18S rRNA metabarcoding versus Western Channel Observatory cell counts by comparison generally showed similar dynamics between the two datasets (**Figure 2B**; **Figure S6A**). However, there were some differences e.g. whereas an increase in the % relative abundance of *Lauderia* was detected in early April by the metabarcoding data (**Figure 2B**), this was not apparent in the cell count data (**Figure S6A**), albeit both datasets did confirm peak *Lauderia* relative abundance in the fourth week of April. Further scrutinising the taxonomic composition of the diatom community and bacterial ASVs through the annual cycle revealed that the early spring (March 2021) samples coinciding with detection of *P. alexandrii* (green line, **Figure 2A**), occurred just prior to a major shift in diatom community in the last week of March, from predominantly *Thalassiosira* to *Chaetoceros* (**Figure 2B**; **Figure S6A**). Late April was characterised by an intense yet brief peak in *Lauderia* (comprising a single species*, Lauderia annulata*) (**Figure 2B; Figure S6A**). Summer months (June 2021 – August 2021) were generally dominated by *Leptocylindrus*, which coincided with the second incidence of *P. alexandrii* (as well as detection of *M. dokdonensis,* alongside *C. atlanticus*) that were detected in the third week of June. Notably, this corresponded with a substantial reduction in *Leptocylindrus* cell counts as detected by the Western Channel Observatory data (from 118 cells/ml to 9 cells/ml) (**Figure S6A**). *V. diazotrophicus* and *Alteromonas* sp. (asv_934) were detected subsequently in the fourth week of July as *Leptocylindrus* populations began to diminish (**Figure 2B**). *Pseudo-nitzschia* species made up the dominant diatom taxa during the large summer diatom bloom in August 2021 (**Figure 2B; Figure S6A**), when at least four bacterial antagonists were observed, including *Halomonas* sp., *M. adhaerens* and *T. lohafexi,* and *Alteromonas sp.* (asv_1961), and when the abundance of bacterial antagonists was at its highest (**Figure 2A**).

Further examination of the species composition of the *Pseudo-nitizschia* ASV population revealed that the August peak was dominated predominantly by ASVs phylogenetically similar to a potentially toxic species *Pseudo-nitzschia fraudulenta* (Tatters, Fu and Hutchins 2012) (**Figure 3A**). However, while the Western Channel Observatory data detected peaks in *Pseudo-nitzschia* abundance in August 2021, the taxonomic assignment (which is limited using light microscopy and therefore only based on cell size and morphological shapes of three distinct groups) differed to the molecular data, with the ‘*P. delicatissima* group’ identified as the most abundant during this time (**Figure S6B**). Plotting total % relative abundance of all antagonistic bacterial ASVs over the annual cycle showed that they closely mirrored *Pseudo-nitzschia* ASVs at three temporally distinct time-points (early June, late August and early November) (**Figure 3A**). Pearson’s Correlation Analysis confirmed a statistically significant positive correlation between total antagonistic bacterial ASV abundances and total *Pseudo-nitzschia* ASV counts (R=0.82; *p*=0.00001) (**Figure 3B**). Additionally, a significant positive correlation was also found between antagonistic bacterial ASV abundances and temperature (R=0.59, *p*=0.006).

**Figure 3.**
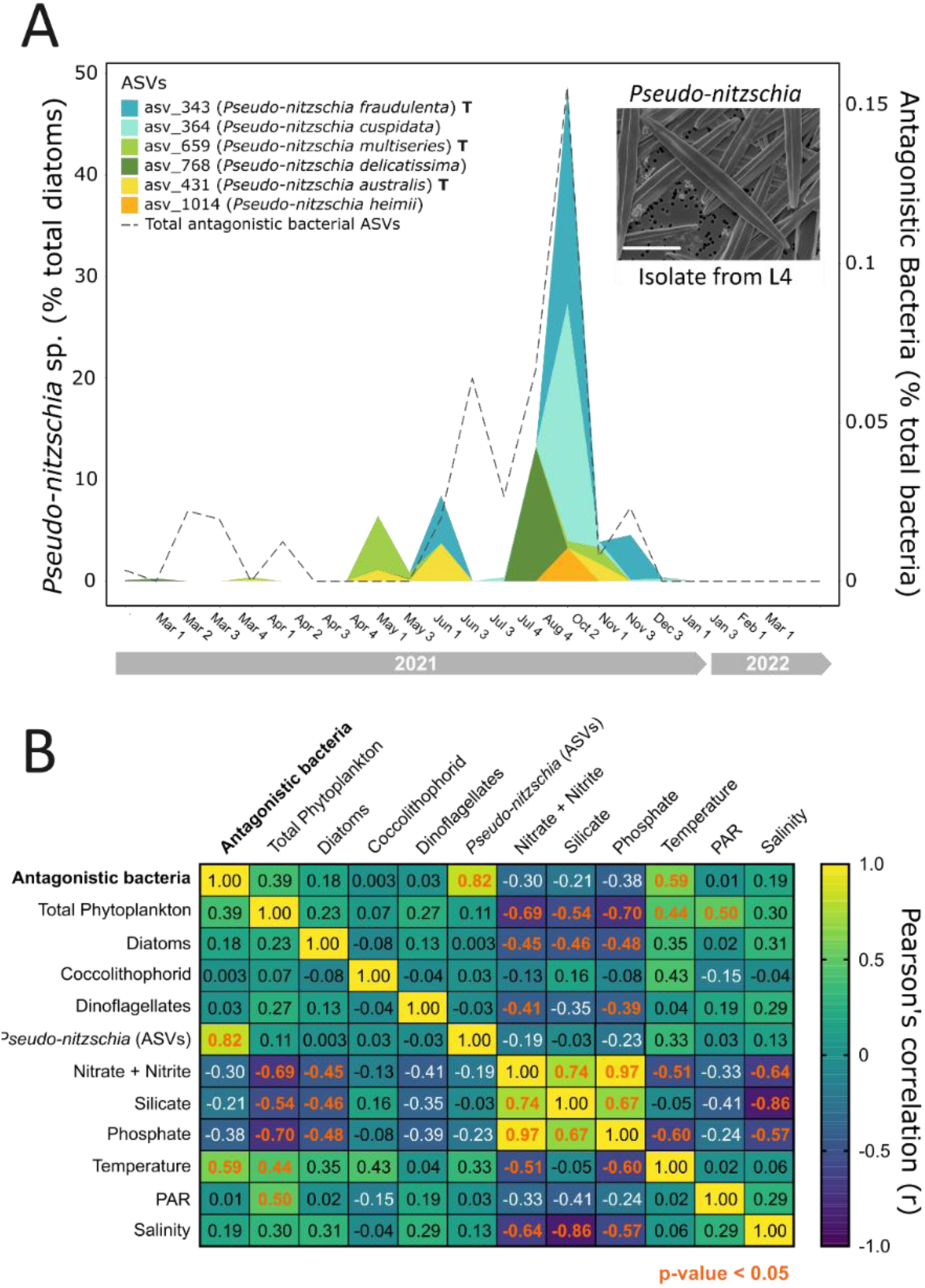
Co-occurrence dynamics of antagonistic bacteria and *Pseudo-nitzschia* at L4 Station. **A.** Percentage relative abundance of total ASVs phylogenetically related to query sequences of bacterial antagonists, and relative abundance of *Pseudo-nitzschia* ASVs as a percentage of total diatom ASVs (left axis) between March 2021 and March 2022 (labelled by sampling week). Species known to produce the harmful toxin domoic acid according to (Tatters et al., 2012; Trainer et al., 2012) are indicated with a ‘T’. Inset shows a representative image of *Pseudo-nitzschia delicatissima* previously isolated from L4 station, imaged via Scanning Electron Microscopy. Scale bar: 10 µm. **B**. Heatmap illustrating a two-way Pearson’s Rank Correlation analysis between total antagonistic bacterial ASV abundance, phytoplankton (including diatom, coccolithophorid and dinoflagellate) abundance (cells/ml), as well as a range of abiotic variables. Total ASVs phylogenetically assigned as *Pseudo-nitzschia* were also included in the analysis. Values between 0 and 1 indicate a positive correlation, whereas values between 0 and -1 show a negative correlation. No relationship between the variables is indicated by ‘0’. Statistically significant correlations (*p* < 0.05) are labelled in orange.

After the summer peaks in diatom abundance the late autumn and winter diatom community was heavily dominated by diatoms belonging to the polar-centric Mediophyceae (order), coinciding with detection of *M. idriensis* in November 2021 (**Figure 2**). No bacterial antagonists were detected from this point onwards. However, the relative abundance of diatoms belonging to the *Coscinodiscus* genus also began to increase between January 2022 – March 2022, peaking in March (**Figure 2B**). *Coscinodiscus* species frequently dominate winter diatom blooms in the WEC (Widdicombe *et al*. 2010), and whilst *P. alexandrii* was isolated previously from a *Coscinodiscus* bloom it was not detected during this time (**Figure 2A**). However, *Coscinodiscus* abundance in January 2022 was low compared to previous years with biomass reaching 3.2 µg C l^-1^ in January 2022 versus 17.8 µg C l^-1^ in December 2020 when *P. alexandrii* was originally isolated (Branscombe *et al*. 2024). Diatoms belonging to the genus *Minidiscus* that also comprises an important part of the diatom community at L4 Station in winter (Arsenieff *et al*. 2020) were also detected both in late March 2021 and November 2021 – March 2022).

### Seasonal abundance of ASVs representing bacterial genera containing antagonistic species

Finally, because different algicidal species can belong to the same bacterial genera (Coyne, Wang and Johnson 2022), we plotted seasonal abundance of all ASVs phylogenetically assigned (via SILVA) to the genus level of the species examined in this study. Taking this approach, we were able to identify additional ASVs beyond the single ASVs already described (**Figure 2A**) for *Vibrio*, *Alteromonas, Marinobacter*, and *Maribacter*, respectively (**Figure 4**). Of these, *Vibrio* was the most diverse genus, with six ASVs identified in total (**Figure 4A**). The most abundant of these was identified as a potential scallop pathogen *V. pectenicida* (Lambert *et al*. 1998), present at the same time as *V. diazotrophicus* in July/August and coinciding with increases in diatom cell density (fourth week of July). Indeed, using Pearson’s Correlation Analysis, we observed a significant correlation between total *Vibrio* ASVs and diatom cell density (R=0.46; *p*=0.041), as well as temperature (R=0.58; *p*=0.008) (**Figure S7**). Multiple *Alteromonas* ASVs were also present (**Figure 4B**), with two additional ASVs identified to those described in Figure 2A. One of these (*Alteromonas* sp. 1) was detected March to May 2021 alongside January 2022 (blue, **Figure 4B**). The remaining *Alteromonas* sp. ASVs were present between July and October 2021. The *Marinobacter* ASV population comprised four ASVs, the ASV phylogenetically similar to the laboratory characterised *M. adhaerens* antagonistic bacterium (asv_989) (**Figure 2A**) (Branscombe *et al*. 2024) represented the most abundant of these (**Figure 4C**). Together *Marinobacter* ASVs significantly correlated with total *Pseudo-nitzschia* ASVs (R=0.78; *p*=0.0004) (**Figure S7**). Finally, we identified just one additional ASV predicted to be a *Maribacter*, which peaked in early spring, but *Maribacter* ASVs were lower in abundance compared to the other genera described (**Figure 4D**).

**Figure 4.**
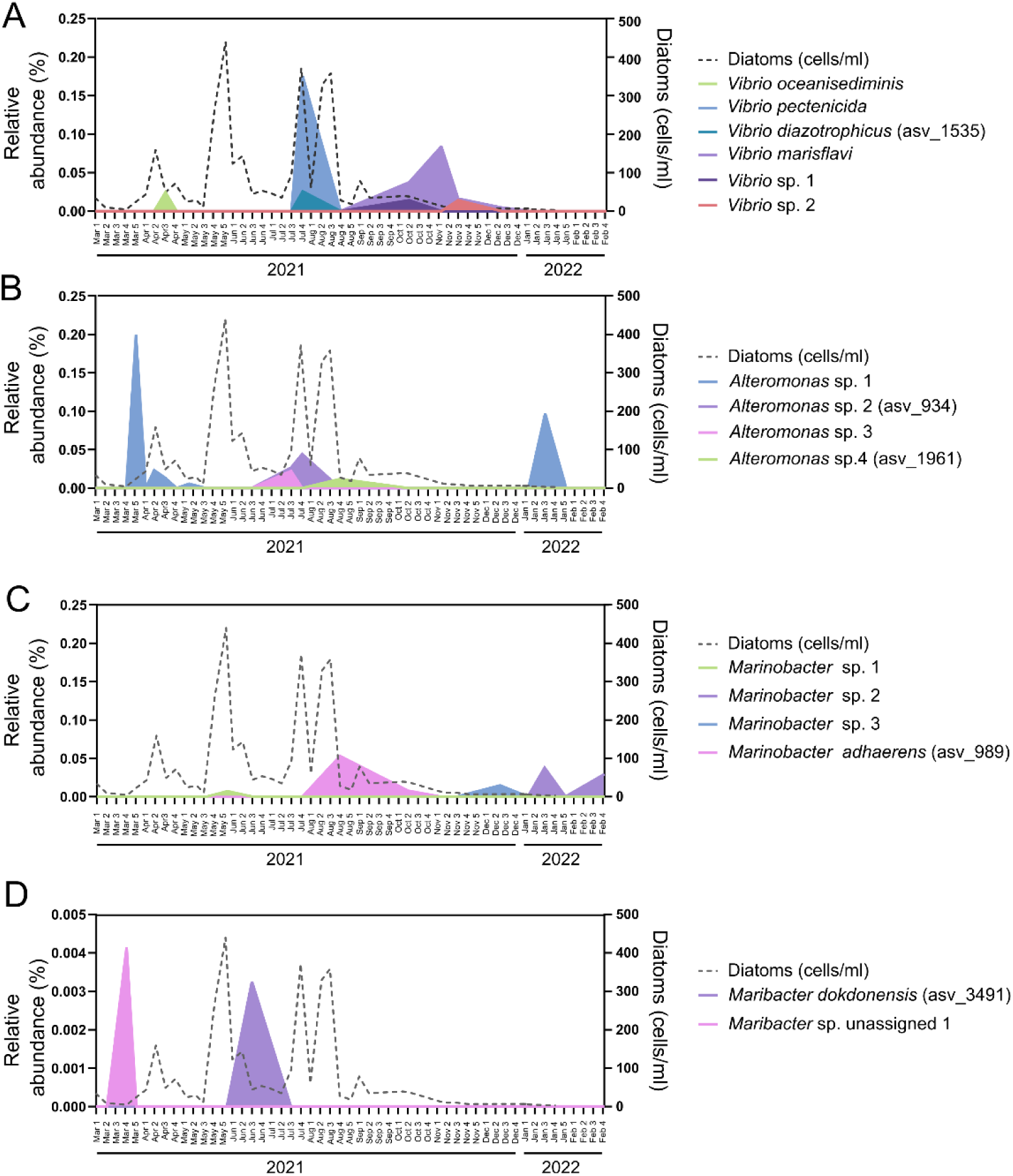
Seasonal abundance of ASVs representing bacterial genera containing antagonistic species. Relative abundance (%) for ASVs phylogenetically assigned to the genera **A.** *Vibrio***, B.** *Alteromonas***, C.** *Marinobacter* and **D.** *Maribacter* (left axis). Total diatom abundance data (cells/ml) are plotted (right axis). For clarity identifiers of ASVs phylogenetically similar to laboratory characterised strains shown in Figure 3 are indicated.

## Discussion

Elucidating the ecological significance of laboratory-characterised diatom-bacteria interactions requires an understanding of their distribution, co-occurrence and seasonal dynamics, and is a fundamental step to better understanding how biotic factors shape marine ecosystem functioning. Techniques such as short-read amplicon sequencing are invaluable for investigating the dynamics of such strains in the natural environment. Utilising these tools, this work has detected and studied ASVs that are phylogenetically similar to multiple facultative bacterial antagonists that, through complementary laboratory work, are known to confer robust growth inhibitory effects against diatoms (Branscombe *et al*. 2024). Further monitoring of these ASVs through the 13-month time series revealed overlapping presence of several antagonists during the summer months (June – August), throughout which the dominant diatom taxa fluctuated from primarily *Leptocylindrus*, to a diverse community of *Thalassiosira*, *Chaetoceros* and *Minidiscus*, to largely *Pseudo-nitzschia*. In addition to studying bacteria previously isolated from this ecosystem, we examined the presence of other algicidal bacteria reported in the literature, including *C. atlanticus* and *A. macleodii* (Van Tol, Amin and Armbrust 2017; Cai *et al*. 2023). Similar to WEC antagonists, ASVs displaying phylogenetic similarity to *C. atlanticus* and *Alteromonas* sp. were detected throughout the summer months (June - August), thus demonstrating that antagonistic bacterial populations in the WEC are highly diverse and exhibit overlapping temporal niches. As well as examining co-occurrence dynamics of antagonistic bacterial populations with dominant eukaryotic phytoplankton taxa, we identified significant positive relationships between facultative antagonistic bacterial (total ASV) abundance and temperature. Our work thus suggests the impacts and abundance of such microbes in the context of future ocean temperatures (van de Waal and Litchman 2020) should be considered.

Consistent with previous literature reports (Widdicombe *et al*. 2010), the diatom communities observed in our study were dynamic, displaying strong seasonal variations in the dominant diatom taxa, where specific genera such as *Leptocylindrus*, *Lauderia*, *Thalassiosira*, and *Pseudo-nitzschia* (amongst others) exhibited seasonal blooms. In many instances we saw occurrence of ASVs phylogenetically similar to bacterial antagonists coincide with shifts in the abundance of diatom species known to be susceptible to the growth inhibitory effects of such bacteria. For example, *P. alexandrii* detection was observed just prior to a decline of *Thalassiosira* in the last week of March (with *Chaetoceros* becoming more dominant) (**Figure 2B**). However, species-specific differences in diatom susceptibility have been observed between members of the same genus (Van Tol, Amin and Armbrust 2017; Meyer and Pohnert 2019; Branscombe *et al*. 2024). The effects of the antagonistic bacteria previously isolated from the WEC on *Pseudo-nitzschia* growth and physiology are not currently known. Nevertheless, *C. atlanticus,* which was originally isolated from an environmental isolate of *Pseudo-nitzschia,* substantially inhibits *P. multiseries* growth by up to 73%, compared to a growth inhibition of only 30% of *P. fraudulenta* AC1 by the same *Croceibacter* strain (Van Tol, Amin and Armbrust 2017). *Pseudo-nitzschia* species are often found abundantly at L4 Station, and frequently bloom during the summer months (Widdicombe *et al*. 2010; Downes-Tettmar *et al*. 2013). The differential susceptibility of different *Pseudo-nitzschia* species to algicidal bacteria could influence *Pseudo-nitzschia* fitness in natural diatom assemblages. Indeed, ASVs phylogenetically assigned as *P. fraudulenta* showed the highest abundance at time points where the greatest number of antagonistic bacterial ASVs were detected. Certain *Pseudo-nitzschia* species (including *P*. *fraudulenta*, *P. multiseries* and *P. australis*) are known producers of the harmful toxin domoic acid (Tatters, Fu and Hutchins 2012; Trainer *et al*. 2012). Low levels of domoic acid have been measured in the WEC, the greatest concentrations correlating with low nitrate and silicate (Downes-Tettmar *et al*. 2013). Domoic acid biosynthesis is well known to be stimulated by nutrient stress (Trainer *et al*. 2012; Brunson *et al*. 2018), however, evidence that bacteria can also enhance *Pseudo-nitzschia* domoic acid production (Bates *et al*. 1995), and are found physically attached to *Pseudo-nitzschia* cells in natural planktonic coastal ecosystems (Kaczmarska *et al*. 2005), is particularly notable in light of our detection of multiple diatom antagonistic bacteria coinciding with *Pseudo-nitzschia* peaks.

Tracking single bacterial ASVs throughout time in a highly dynamic microbial community, consisting of thousands of species, is inevitably challenging, particularly as short-read amplicon sequencing, while a powerful tool for the exploration of microbial communities at an ecosystem scale, is not without limitations or biases. For example, one of the most prominent issues faced with short-read amplicon sequencing pipelines is the taxonomic assignment of ASVs, particularly at genus or species level. Short-read amplicon sequencing utilises subregions of the 16S rRNA gene, termed variable regions, to distinguish separate sequences and assign taxonomy (Ghyselinck *et al*. 2013; Abellan-Schneyder *et al*. 2021). Numerous studies have demonstrated the effects of the chosen variable region used for amplicon sequencing, with some regions better able to discern species than others, thus producing more robust taxonomic assignment (Bukin *et al*. 2019; Callahan *et al*. 2021). Our study utilised the V4 region, widely used for short-read amplicon sequencing as it is the region of the 16S rRNA gene which contains maximum nucleotide heterogeneity (Ghyselinck *et al*. 2013; Yang, Wang and Qian 2016). Nevertheless, multiple studies have reported lower diversity using this variable region compared to other longer regions, for example, the V2-V3 or V3-V4 regions, or even full-length 16S rRNA sequencing. These longer regions allow for a more robust separation of closely related sequences, hence increasing and better representing the true diversity of microbial communities (Bukin *et al*. 2019; Greay *et al*. 2019). As such, it is possible that some diversity of the prokaryotic community was lost throughout the sampling pipeline. Despite these challenges, we successfully detected ASVs phylogenetically similar to nine bacterial antagonists queried that have confirmed growth inhibitory effects towards diatoms (*M. adhaerens*, *M. dokdonensis*, *M. idriensis*, *P. alexandrii*, *T. lohafexi*, *V. diazotrophicus, Halomonas* sp.*, A. simiduii,* and *C. atlanticus*) at nearly half of the timepoints sampled.

While at first glance the relative abundance of the bacterial ASVs that were detected in our study appear low (<1%), many of the species examined (*including P. alexandrii*, *M. adhaerens*, *A. macleodii*, *C. atlanticus* and *Vibrio* sp.) are particle/cell-attaching bacteria (Gärdes *et al*. 2010; Van Tol, Amin and Armbrust 2017; Hubert and Michell 2020; Cai *et al*. 2023; Branscombe *et al*. 2024). Particle-attached bacteria typically represent a lower proportion (0.1-4%) of bacterioplankton abundance during phytoplankton blooms compared to free-living representatives (Alldredge, Cole and Caron 1986; Wang *et al*. 2024). Yet, despite their lower abundance, they are estimated to mediate around 50% of particulate organic matter fluxes in the ocean (Giering *et al*. 2014). Moreover, the generally larger genomes of particle-attached bacteria encode expanded metabolic repertoires for solubilisation and remineralisation of algal-derived polysaccharides (Wang *et al*. 2024). Hence, particle-attaching bacteria have been described as ‘gatekeepers’ in the decomposition of organic matter in planktonic marine communities. Furthermore, in addition to identifying single ASVs phylogenetically similar to facultative antagonistic bacteria characterised in the laboratory (Branscombe *et al*. 2024), we identified additional ASVs assigned to the same genera. Most notably, identifying a significant positive correlation between *Vibrio* relative abundance and diatom cell counts. *Vibrio* species are capable of attaching to diatom cells (Hubert and Michell 2020), and adhering to diatom-derived chitin via type IV pili (Frischkorn, Stojanovski and Paranjpye). Evidence that different *Vibrio* species can also significantly impact diatom growth (Ismail and Ibrahim 2017; Branscombe *et al*. 2024) indicates the need to further explore *Vibrio*-diatom interactions.

Whilst in general we observed good consensus between molecular sequencing and the eukaryotic phytoplankton community abundance data obtained via Western Channel Observatory cell counts, there were a few notable exceptions. Firstly, whereas the 18S rRNA gene data determined dinoflagellates to be generally more abundant than diatoms at L4 Station (**Figure S5**), the reverse was observed for the cell count data (**Figure S1**). Dinoflagellates can have very high rDNA gene copy numbers (Ruvindy et al. 2023), which likely explains this discrepancy. In contrast 18S rRNA gene copy number does not seem to be a major concern for diatoms (Malviya *et al*. 2016). However, even though genus-level trends in the abundance of *Pseudo-nitzschia* held true between our two datasets, there were differences in the species level assignments. Whereas the 18S rRNA gene amplicon sequencing data identified *P. fraudulenta* as the most abundant taxa during the bloom peaking in August 2021 (**Figure 3A**), the Western Channel Observatory cell count data identified this peak to comprise mainly the ‘*P. delicatissima* group’, based on cell size and morphology determined by light microscopy (Downes-Tettmar *et al*. 2013). *Pseudo-nitzschia* are notoriously challenging to identify to species level based on morphological shape without detailed analysis using electron microscopy (Lundholm, Daugbjerg and Moestrup 2002; Lundholm *et al*. 2006). Similarly, the 18S rRNA V9 region (used in this study) is not ideal for distinguishing diatom species (Malviya *et al*. 2016). As such the phylogenetic assignment of *Pseudo-nitzschia* to species level should be taken with caution.

Finally, our study has focused on bulk sampling of diatom and bacterial populations. Whilst this approach is informative in assessing overarching trends in presence and co-occurrence of specific diatoms and bacterial taxa, clearly there is a need to study diatom-bacteria interactions in natural planktonic ecosystems on the microscale. Several of the bacteria queried in this study (e.g. *M. adhaerens* and *P. alexandrii*) are capable of physically attaching to diatoms (Gärdes *et al*. 2010; Branscombe *et al*. 2024). Indeed, *M. adhaerens* is also chemotactic towards diatom cells, attracted to diatom exudates (Sonnenschein *et al*. 2012). Marine ecosystems are incredibly varied and dilute, however under certain circumstances the conditions may promote antagonistic interactions between diatoms and bacteria. Intense spring blooms exceeding 4000 cells ml^-1^ of diatoms such as *Chaetoceros* sp. have been reported in the WEC (Widdicombe *et al*. 2010). Cell densities of this magnitude, combined with bacterial pathogen chemotaxis and motility, would enhance encounter rates between diatoms and bacteria (Seymour *et al*. 2017), and likely exacerbate potential impacts of algicidal bacteria on diatom productivity. Work using single-cell environmental isolates of *Thalassiosira rotula* have provided important insights of how host genotype and biogeography may impact diatom microbiome structure (Ahern *et al*. 2021). Efforts must now focus on studying temporal patterns in the diversity and function of the bacterial communities and diatoms at the single-cell level over the course of diatom bloom succession. Such studies will be facilitated by advances in our understanding of the genes underlying algicidal behaviour (Syhapanha *et al*. 2023), which will provide bioindicators necessary to quantify algicidal activity in the field, and is especially important given the often facultative pathogenicity observed amongst algicidal bacteria (Seyedsayamdost *et al*. 2011; Barak-Gavish *et al*. 2023).

## Acknowledgements

We are thankful for support from the NERC Independent Research Fellowship grant NE/R015449/2 (K.E.H.) and PhD studentships from the NERC ARIES doctoral training program NE/S007334/1 (L.B. and C. R.). C.E.W. was supported by funding from the NERC National Capability Long-term Single Centre Science Programme, Climate Linked Atlantic Sector Science, grant number NE/R015953/1. We also thank crew of the RV Sepia (MBA, Plymouth, UK) and Plymouth Quest (PML, Plymouth, UK) for their collection of samples used during this study. Glen Tarran (PML, Plymouth, UK) and Malcolm Woodward (PML, Plymouth, UK) kindly shared the Station L4 flow cytometry and nutrients data, respectively.

## Data availability

Data will be deposited into the European Nucleotide Archive (ENA) upon acceptance for publication.

## Notes

### Competing Interest Statement

The authors have declared no competing interest.

