## Supplementary Information for "Seasonal niche overlap of diverse facultative antagonistic bacteria of diatoms in a productive coastal ecosystem"

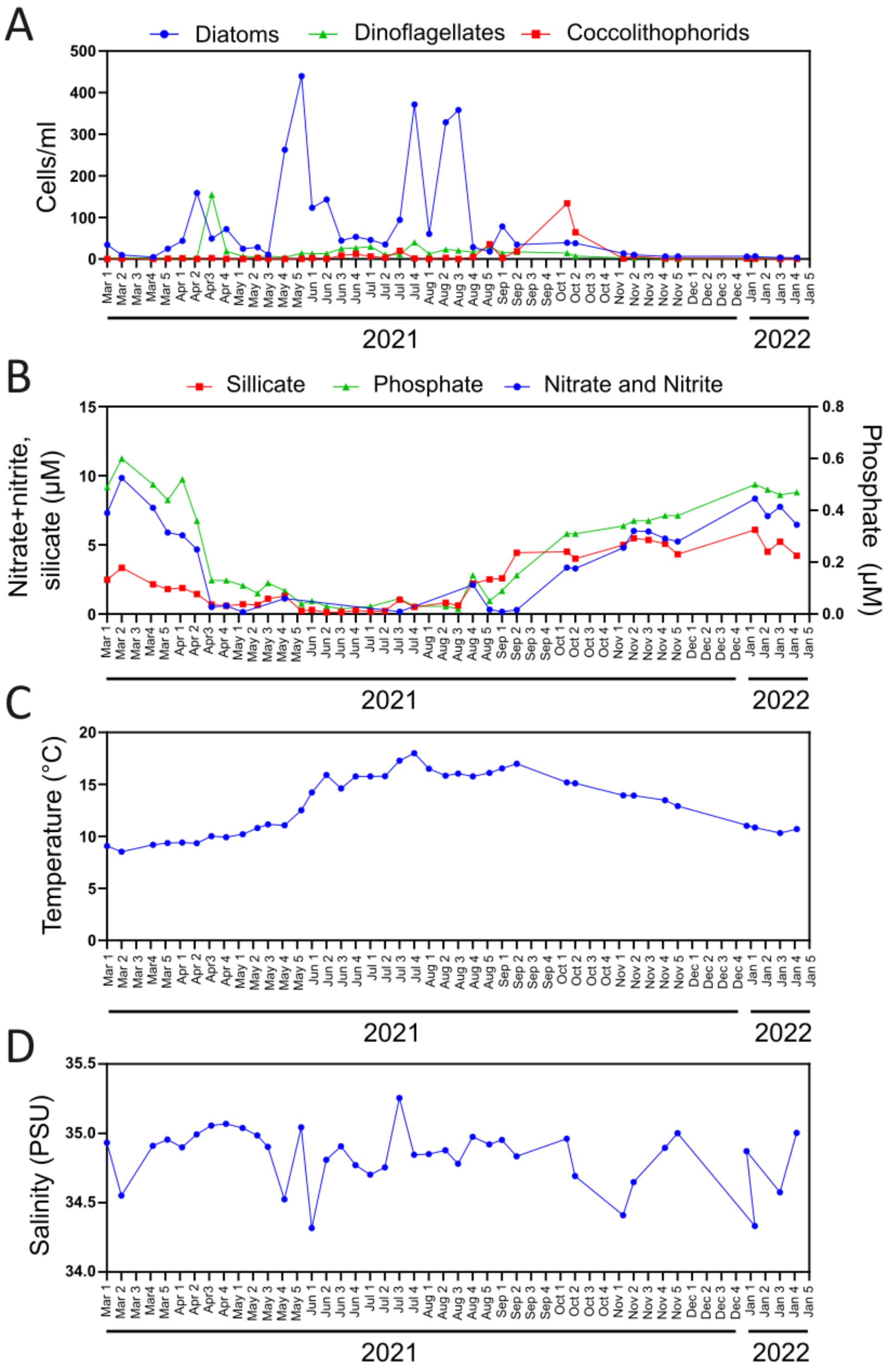


**Figure S1 Environmental metadata collected at L4 Station over the sampling period. A.** Abundance (cells/ml) of major phytoplankton classes: diatoms, coccolithophorids and dinoflagellates. **B.** Concentrations (µM) of macronutrients silicate, nitrate and nitrite as well as phosphate, at 10 m depth. Data for temperature (°C) (**C**), and salinity (practical salinity unit, PSU) at 5 m depth (**D**) are also plotted. All data are from the Western Channel Observatory**.**

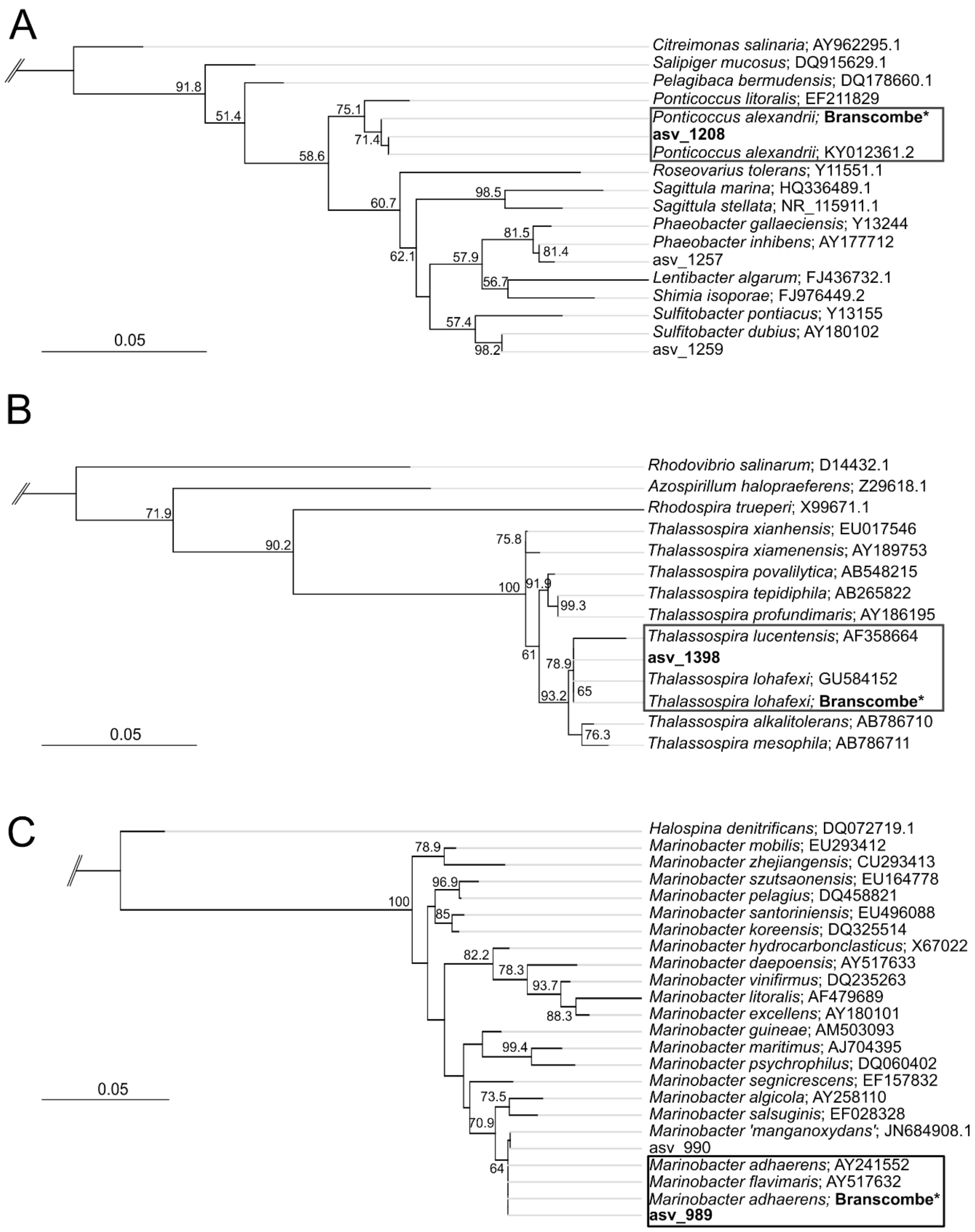


**Figure S2. Maximum Likelihood trees verifying the identity of amplicon sequence dataset ASVs, including**: **A.** *Ponticoccus alexandrii.* **B.** *Thalassospira lohafexi*. and **C.** *Marinobacter adhaerens*, using *Rubellimicrobium aerolatum*, *Azorhizobium caulinodans* and *Salicola marasensis* as outgroups for each tree, respectively. Maximum Likelihood trees were constructed using ASV sequences as well as 16S rRNA sequences of query bacteria (bacteria with confirmed antagonistic activity from (Branscombe et al., 2024), are labelled “Branscombe*”), and type strains of closely related species. Branch support values above 50% are indicated at branch nodes (calculated from 1000 bootstrap replicates).

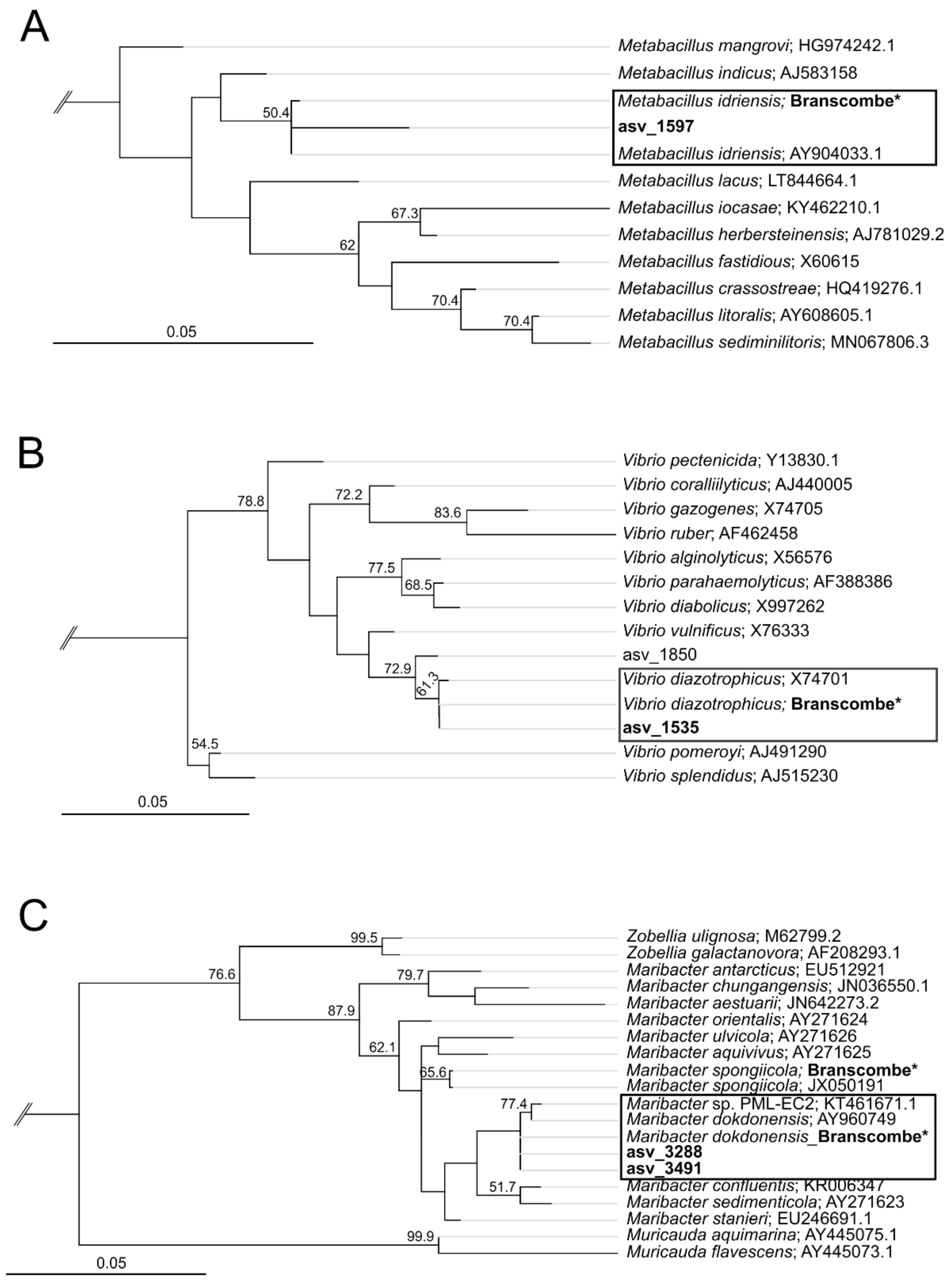


**Figure S3 Maximum Likelihood trees verifying the identity of amplicon sequence dataset ASVs, including**: **A.** *Metabacillus idriensis*. **B.** *Vibrio diazotrophicus* and **C.** *Maribacter dokdonensis,* using *Lysinibacillus boronitolerans, Salinivibrio costicola* and *Flavobacterium aquatile* as outgroups for each tree, respectively. *Maribacter spongiicola* is also included. Maximum Likelihood trees were constructed using ASV sequences as well as 16S rRNA sequences of query bacteria (bacteria with confirmed antagonistic activity from (Branscombe et al., 2024), labelled 'Branscombe*'), and type strains of closely related bacterial species. Branch support values above 50% are shown (calculated from 1000 bootstraps).

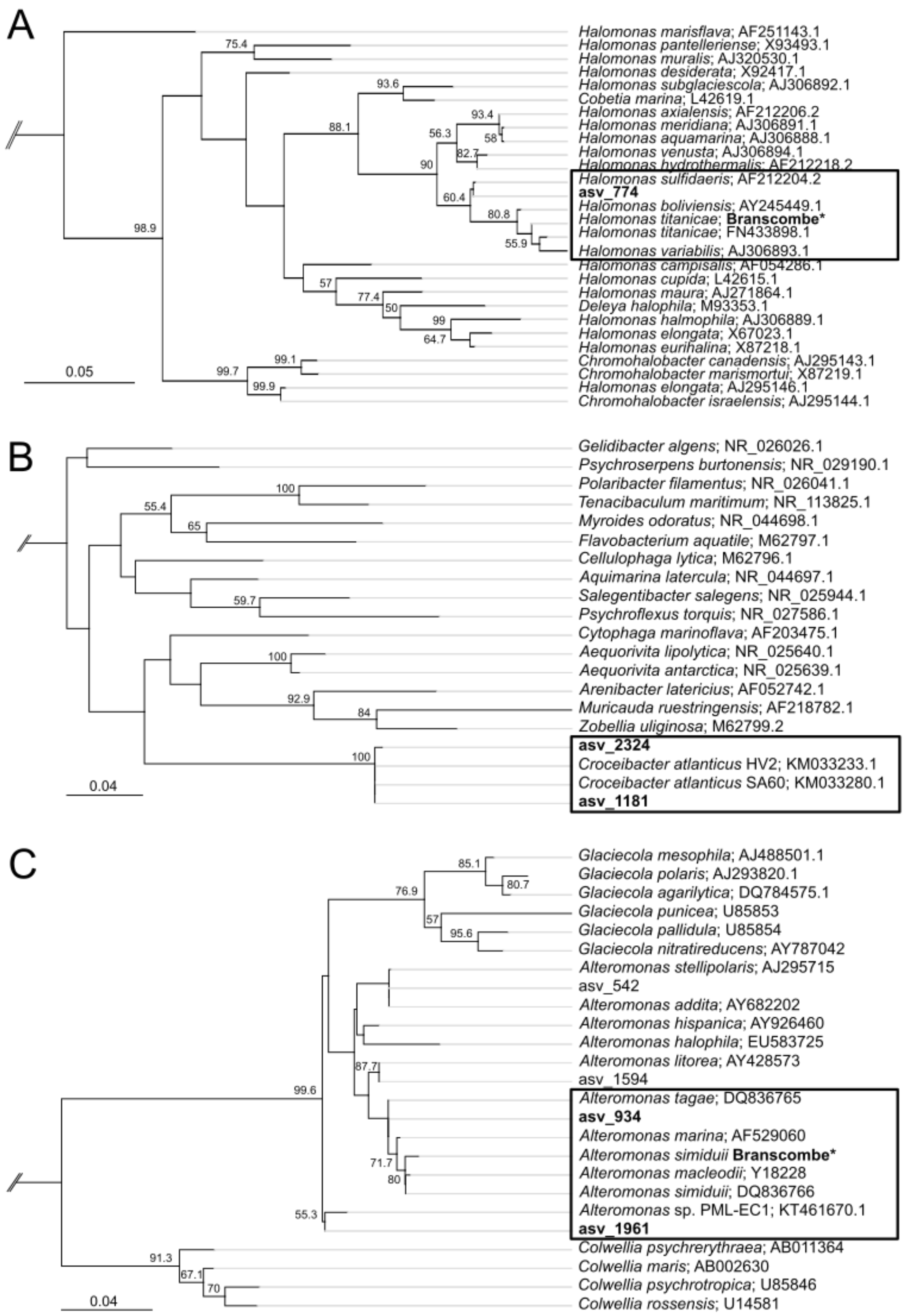


**Figure S4. Maximum Likelihood trees constructed to verify the identity of amplicon sequence dataset ASVs most closely related to A.** *Halomonas titanicae***, B.** *Croceibacter atlanticus* and **C.** *Alteromonas* sp., using *Zymobacter palmae, Pseudomonas aeruginosa* and *Coenonia anatina* as outgroups for each tree, respectively. Maximum Likelihood trees were constructed using ASV sequences as well as 16S rRNA sequences of query bacteria (bacteria with confirmed antagonistic activity from (Branscombe et al., 2024), are labelled 'Branscombe*', where applicable), and type strains of closely related bacterial species. Branch support values above 50 % are shown (calculated from 1000 bootstraps).

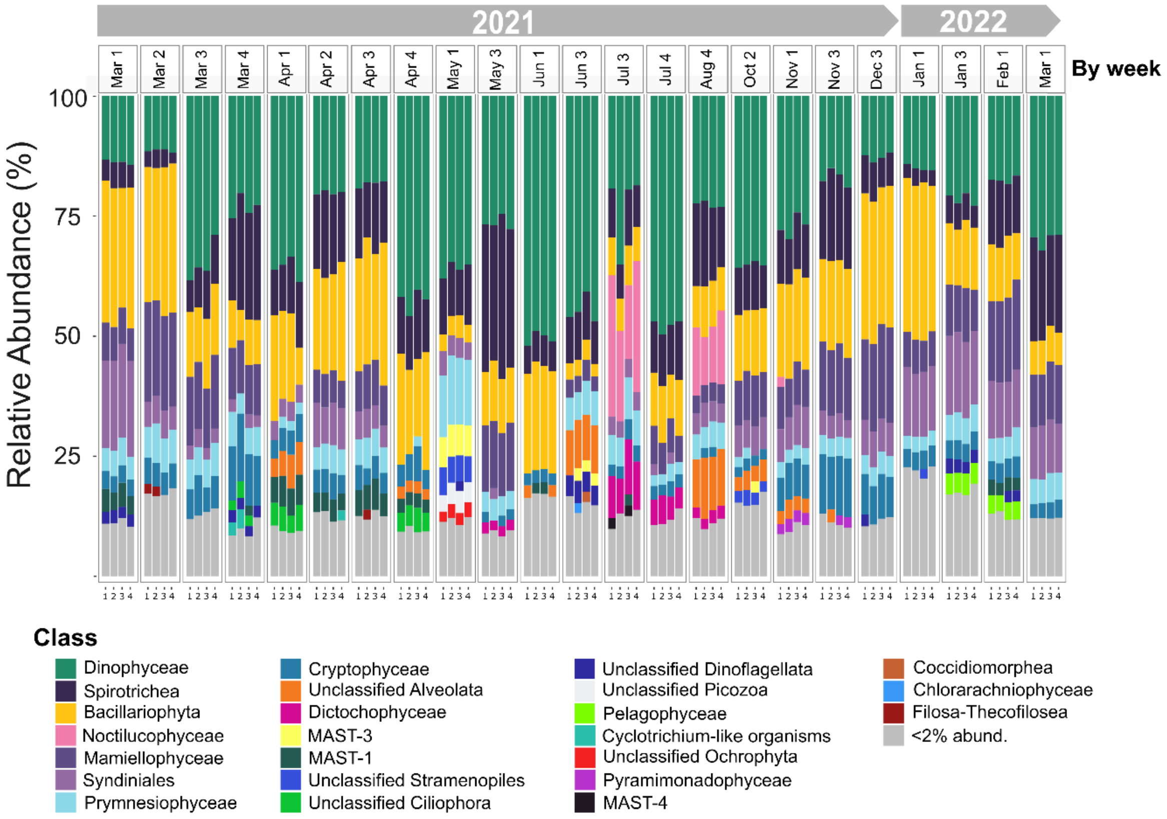


**Figure S5. Percentage relative abundance of amplicon sequencing variants (ASVs) belonging to major phytoplankton classes over an annual cycle.** Highlighting the seasonal persistence and abundance of diatoms (Bacillariophyta, yellow). Bars represent the four replicates processed at each sampling point. Taxa that constituted less than 2% of the total ASVs are grouped into the <2% abundance category.

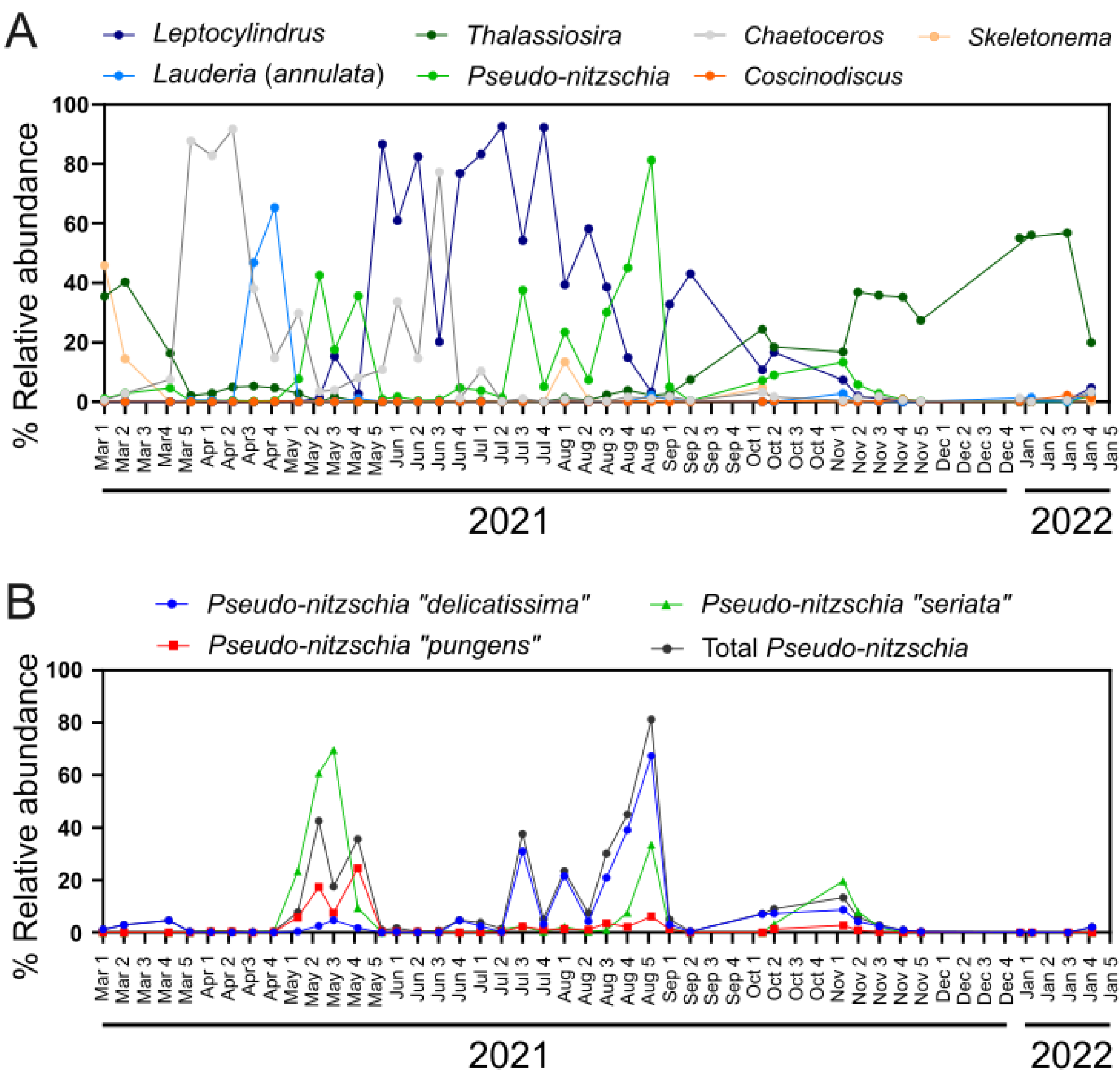


**Figure S6. Trends in diatom abundance as determined by Western Channel Observatory cell counts. A.** Relative abundance (as a % of total diatoms) of seven major diatom genera between March 2021 and Jan 2022. **B.** Relative abundance (as a % of total diatoms) of three *Pseudo-nitzschia* groups (*P. delicatissima*, *P. pungens* and *P. seriata*) phylogenetically assigned according to morphology and cell size by light microscopy (Downes-Tettmar et al., 2013; Widdicombe et al., 2010).

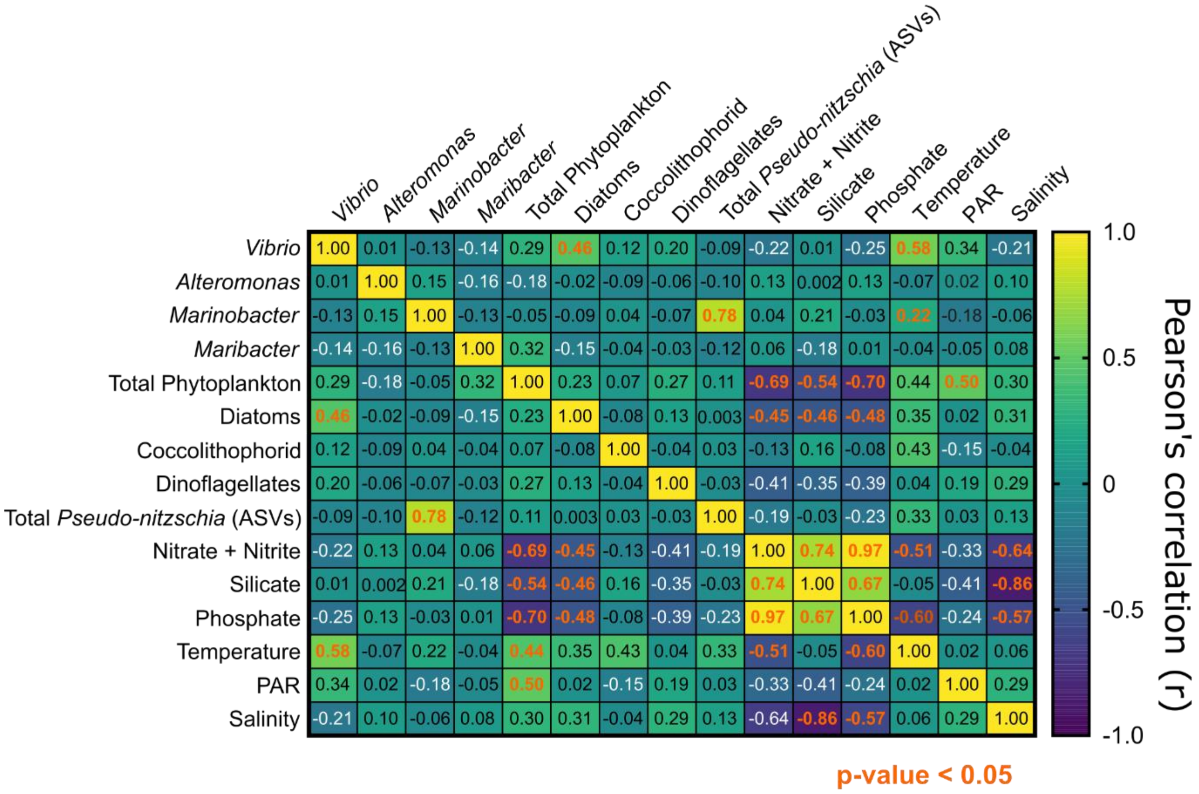


**Figure S7**. **Pearson's correlation analysis of antagonistic bacterial genera and abiotic and biotic variables**. Heatmap illustrating a two-way Pearson's Rank Correlation analysis between total ASVs for *Vibrio*, *Alteromonas*, *Marinobacter*, *Maribacter* as well as phytoplankton (including diatom, coccolithophorid and dinoflagellate) abundance (cells/ml), and a range of abiotic variables. Total ASVs phylogenetically assigned as *Pseudo-nitzschia* were also included in the analysis. Values between 0 and 1 indicate a positive correlation, whereas values between 0 and -1 show a negative correlation. No relationship between the variables is indicated by '0'. Statistically significant correlations ($p$ < 0.05) are labelled in orange

**Table S1.** List of sampling dates for metabarcoding samples versus the Western Channel Observatory (WCO) data.

| Metabarcoding sampling date | WCO sampling date |
|---|---|
| 01-Mar-2021 | 01-Mar-21 |
| 09-Mar-2021 | 08-Mar-21 |
| 23-Mar-2021 | 23-Mar-21 |
| 30-Mar-2021 | 30-Mar-21 |
| 06-Apr-2021 | 06-Apr-21 |
| 12-Apr-2021 | 13-Apr-21 |
| 20-Apr-2021 | 20-Apr-21 |
| 28-Apr-2021 | 27-Apr-21 |
| 07-May-2021 | 05-May-21 |
| - | 12-May-21 |
| 19-May-2021 | 17-May-21 |
| - | 25-May-21 |
| 03-Jun-2021 | 02-Jun-21 |
| - | 07-Jun-21 |
| - | 14-Jun-21 |
| 21-Jun-2021 | 21-Jun-21 |
| - | 28-Jun-21 |
| - | 05-Jul-21 |
| - | 12-Jul-21 |
| 19-Jul-2021 | 19-Jul-21 |
| 27-Jul-2021 | 26-Jul-21 |
| - | 02-Aug-21 |
| - | 10-Aug-21 |
| - | 16-Aug-21 |
| 23-Aug-2021 | 23-Aug-21 |
| - | 31-Aug-21 |
| - | 06-Sep-21 |
| - | 13-Sep-21 |
| - | 07-Oct-21 |
| 11-Oct-2021 | 11-Oct-21 |
| 03-Nov-2021 | 03-Nov-21 |
| - | 08-Nov-21 |
| 15-Nov-2021 | 15-Nov-21 |
| - | 23-Nov-21 |
| - | 29-Nov-21 |
| 16-Dec-2021 | - |
| 01-Jan-2022 | 01-Jan-22 |
| 17-Jan-2022 | 17-Jan-22 |
| - | 25-Jan-22 |
| 01-Feb-2022 | - |
| 01-Mar-2022 | - |

**Table S2. Samples removed from the 16S rRNA amplicon sequence dataset due to low read depth to reduce diversity loss.** Sample date, replicate number (four replicates were processed per sampling point) and read depth of each sample are given.

| Sample | Sampling Date | Replicate | Sampling Location | Read depth |
|---|---|---|---|---|
| P1KBird41 | 30/03/2021 | 1 | Station L4 | 8,391 |
| P1KBird42 | 30/03/2021 | 2 | Station L4 | 6,753 |
| P1KBird73 | 28/04/2021 | 1 | Station L4 | 8,966 |
| P1KBird75 | 28/04/2021 | 3 | Station L4 | 6,043 |

**Table S3. Top amplicon sequencing variant (ASV) hits for each bacterial antagonist.** Sequences were queried against the raw (non-rarefied) 16S amplicon sequence dataset. ASV number, percentage pairwise identity when aligned with query sequence, and the bacterial sequence that an ASV most closely aligns with under Maximum Likelihood tree construction are also provided. Percentage pairwise identity covers a 254 base pair fragment as this was the maximum length of ASVs in the amplicon sequence dataset.

| **Bacterial antagonist** | **Amplicon sequence ASV hits** | **% Pairwise identity** | **Phylogenetic grouping inferred from Maximum Likelihood trees** |
|---|---|---|---|
| *Ponticoccus alexandrii* | asv_1208 | 100 | *P. alexandrii* |
| | asv_1257 | 98.8 | *Phaeobacter inhibens* |
| | asv_1259 | 97.6 | *Sulfitobacter dubius* |
| *Thalassospira lohafexi* | asv_1398 | 100 | *T. lohafexi* |
| *Marinobacter adhaerens* | asv_989 | 100 | *M. adhaerens* |
| | asv_990 | 99.6 | *Marinobacter 'manganoxydans'* |
| *Metabacillus idriensis* | asv_418 | 99.3 | *Metabacillus mangrovi* |
| | asv_1597 | 98.5 | *M. idriensis* |
| *Vibrio diazotrophicus* | asv_1535 | 100 | *V. diazotrophicus* |
| | asv_1850 | 99.2 | Unresolved |
| *Maribacter spongiicola* | asv_3491 | 97.6 | *M. dokdonensis* |
| | asv_3288 | 95.7 | *M. dokdonensis* |
| *Halomonas titanicae* | asv_774 | 100 | *H. sulfidaeris* |
| *Croceibacter atlanticus* | asv_1181 | 100 | *C. atlanticus* |
| | asv_2324 | 99.6 | *C. atlanticus* |
| *Alteromonas macleodii* | asv_934 | 100 | *Alteromonas tagae* |
| | asv_1961 | 99.6 | *Alteromonas* sp. PML-EC1 |
